# Oligodendrocyte stress response as a driver of vascular dementia

**DOI:** 10.64898/2026.10.12.758545

**Authors:** Emeric Merour, Trupti Ghatage, Cole Corbett, Mathew DeBerge, Rejani Kunjamma, Ian Steckler, Moones Heidari, Farzaneh A. Sorond, Margaret E. Flanagan, Edward B. Thorp, Brian Popko

## Abstract

Vascular dementia (VaD) is a prevalent and devastating neurocognitive disorder characterized by profound white matter damage, yet the precise cellular mechanisms linking cerebrovascular stress to demyelination remain poorly understood. Here, we characterize the impact of a chronic cardiometabolic stress regimen combining a high-fat diet and hypertension (HF/HBP) on central nervous system structure and function. Adult HF/HBP mice exhibit progressive oligodendroglial cell loss and myelin thinning, followed by cognitive impairments, anxiety-like behavior, and memory deficits that recapitulate clinical features of VaD. Mechanistically, we identify the integrated stress response (ISR) as a central driver of this pathology; pharmacological inhibition of the ISR restores myelination, while cell-specific genetic deletion of the ISR-activating kinase PERK from oligodendrocytes fully protects mice from HF/HBP-induced pathological and behavioral defects. Furthermore, we provide translational evidence that the ISR is activated in oligodendrocytes in human cases of cerebrovascular disease. Together, these findings reveal a critical, cell-autonomous role for oligodendroglial ISR activation during cerebrovascular stress and position this pathway as a viable therapeutic target to mitigate VaD pathogenesis.

## Main

Dementia currently affects over 55 million people worldwide, a figure projected to triple by 2050^1^. Clinically, dementia is characterized by significant memory loss alongside the impairment of at least one other cognitive function, such as aphasia or agnosia^2^. This cognitive decline is frequently accompanied by neuropsychiatric symptoms, including apathy, depression, and anxiety^3^. Although Alzheimer’s disease represents the predominant form of dementia, vascular dementia (VaD), driven by cardiovascular pathophysiology, accounts for up to twenty percent of dementia cases. VaD is characterized by the appearance of white matter hyperintensities (WMH) detected by magnetic resonance imaging (MRI)^4,5^. WMH are areas of demyelination and serve as one of the most reliable biomarkers for neurovascular cognitive decline^5^. Despite its massive global burden, VaD research is impeded by a lack of robust preclinical models to test disease-modifying therapies.

The development of a representative preclinical model for VaD requires recapitulating its primary clinical drivers, namely obesity^6,7^ and chronic hypertension^8^. Traditional experimental models, most notably Bilateral Common Carotid Artery Stenosis, simulate vascular cognitive impairment through abrupt and severe surgical constriction of cerebral blood flow^9^. Nevertheless, these models cause an acute ischemic profile that is too severe to accurately represent the long-term, systemic microvascular disease seen in the majority of human VaD cases. To identify a more physiologically accurate approach, we turned to a dual-hit metabolic and hypertensive protocol originally described by Schiattarella et al.^10^. By pairing a high-fat diet with the nitric oxide synthase antagonist N[w]-nitro-l-arginine methyl ester (L-NAME), they modeled human heart failure with preserved ejection fraction, which encompasses over half of all heart failure cases. Crucially, these dietary and chemical interventions induce profound systemic weight gain and elevated blood pressure without disrupting cardiac ejection fraction. In this study, we utilize this high-fat/high-blood-pressure (HF/HBP) model to establish a more translationally relevant baseline for investigating VaD pathology and potential therapeutics.

To evaluate the translational relevance of this non-surgical HF/HBP paradigm, we tracked its progressive impact on the CNS over a 10-weeks period. After 5 weeks of HF/HBP, we detected localized white matter degeneration, specifically characterized by the depletion of oligodendrocytes and structural thinning of myelin sheets. By week 10, this underlying neuropathology manifested behaviorally as multifaceted cognitive decline, including impaired object recognition, heightened anxiety, and learning deficits that mirror the clinical profile of human neurovascular disease. Investigating the molecular drivers behind this degeneration, we identified the integrate stress response (ISR) as a central mediator of vascular-induced CNS damage. Strikingly, pharmacological inhibition of this pathway ameliorated the adverse effects of the HF/HBP regimen. Similarly, genetic ablation of the upstream ISR kinase PERK specifically in oligodendrocytes shielded the HF/HBP mice from myelin damage and subsequent behavioral decline. By confirming that this same cellular stress pathway is actively upregulated in human VaD brain tissue, our data not only validate the HF/HBP protocol as a highly predictive model but also highlight ISR inhibition as a promising therapeutic strategy to halt disease progression.

## RESULTS

### Prolonged HF/HBP Diet Induces Multifaceted Cognitive Deficits

To validate the translational utility of the HF/HBP paradigm for modeling VaD, we evaluated the timeline of cognitive decline in adult mice. Behavioral cohorts were assessed at 5 weeks—the earliest time point previously linked to cardiac phenotypes^10^— or after 10 weeks of diet to track disease progression (**Fig. 1a**). Accompanying their cardiovascular stress, which we previously validated^11^, HF/HBP mice exhibited rapid and progressive systemic weight gain, outpacing control animals by an average of 6g at 5 weeks and 14g by 10 weeks (**Fig. 1b**).

**Figure 1.**
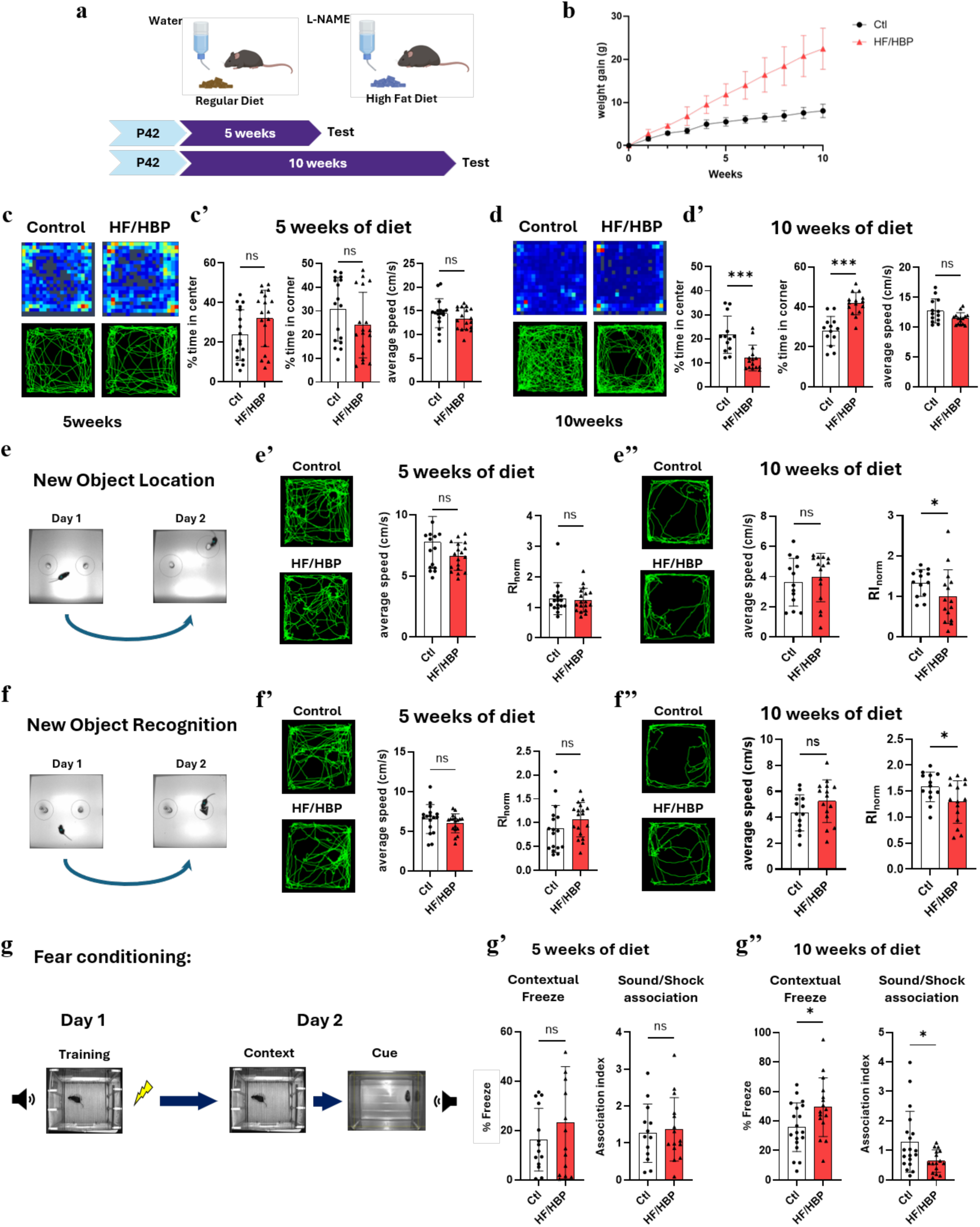
Cognitive Impairment in Mice Following 10 Weeks of HF/HBP Diet. (**a**) Timeline of the experiment. (**b**) Weekly weight-gain in WT Control and HF/HBP male mice (n=20 Control (Ctl), 20 HF/HBP, Area under curve analysis (AUC): Control area = 49.40, 95% Confidence Interval: 45.25 to 53.55; HF/HBP area = 116.3, 95% Confidence Interval: 102.7 to 129.8). (**c**) Open Field (OF) test at 5 weeks of diet with representative path maps and position heatmaps. (**c’**) Percentage of time in the center or corners of the box and average speed (n=17 Ctl, 18 HF/HBP, Mann-Whitney: “% of time in center” p = 0.077; “% of time in corner” p = 0.219; average speed p = 0.443). (**d**) OF at 10 weeks with representative path maps and position heatmaps. (**d’**) Percentage of time in the center or the corners of the box and average speed (n=13 Ctl, 15 HF/HBP, Mann-Whitney: “% of time in center” p < 0.001; “% of time in corner” p < 0.001; average speed p = 0.080). (**e**) Novel Object Location (NOL) test with representative path maps. (**e’**) Speed and RI_norm_ at 5 weeks (n=17 Ctl, 18 HF/HBP, Mann-Whitney: RI_norm_ P = 0.987; “average speed”, P = 0.083) (**e’’**). Speed and RI_norm_ at 10 weeks (n=13 Ctl, 15 HF/HBP, Mann-Whitney: RI_norm_ p = 0.046; “average speed” p = 0.142). (**f**) New Object Recognition (NOR) test with representative Path maps. (**f’**) Speed and RI_norm_ at 5 weeks (n=17 Ctl, 18 HF/HBP, Mann-Whitney: RI_norm_ p = 0.072; “average speed” p = 0.163) (**f’’**). Speed and RI_norm_ at 10 weeks (n=13 Ctl, 15 HF/HBP, Mann-Whitney: RI_norm_ p = 0.041; “average speed” p = 0.586). (**g**) Fear Conditioning (FC) test following 5 and 10 weeks of diet. (**g’**) Percentage of freezing during the contextual test and Association Index (AI) during the cued assay at 5 weeks (n=13 Ctl, 15 HF/HBP, Mann-Whitney: “Contextual Freeze” p = 0.301; “AI” p = 0 .446) (**g’’**). Percentage of freezing during the contextual test and AI during the cued assay at 10 week (n=19 Ctl, 16 HF/HBP, Mann-Whitney: “Contextual Freeze” p = 0.026; “AI” p = 0.048)

To decouple cognitive performance from gross motor function and baseline anxiety-like behavior, mice were first evaluated using the Open Field (OF) test. At the 5-week mark, exploratory drive, total locomotion, and zone preferences were indistinguishable between the experimental and control groups (**Fig. 1c,c’**). However, by 10 weeks, HF/HBP mice displayed an anxiety-like phenotype and altered motor patterns, characterized by a decreased time spent exploring the center zone and a corresponding increase in thigmotaxis (corner-dwelling time) with no change in locomotion (average speed) (**Fig. 1d,d’**).

We next examined spatial memory and object recognition using the novel object location (NOL) and novel object recognition (NOR) paradigms. At 5 weeks, both groups exhibited normal spatial working memory, spending significantly more time exploring the displaced object during the NOL test (**Fig. 1e–e’**). By 10 weeks, this preference was entirely abolished in the HF/HBP cohort; their normalized recognition index (RI_norm_) hovered near 1.0, signifying a failure to detect the spatial shift (**Fig. 1e’’**). Similarly, object recognition remained intact at 5 weeks but was severely impaired at 10 weeks, as evidenced by a marked reduction in the RI_norm_ of the HF/HBP mice when presented with a novel object (**Fig. 1f–f’’**). Notably, mean velocities during the NOL and NOR tasks did not differ between cohorts at either time point, confirming that these recognition deficits were not confounded by physical lethargy or localized locomotor bias.

Finally, associative learning and memory were challenged using a pavlovian fear-conditioning (FC) paradigm (**Fig. 1g**). Consistent with our recognition data, no differences in freezing behavior were observed at 5 weeks (**Fig. 1g’**). However, at 10 weeks, HF/HBP mice demonstrated distinct associative deficits during the contextual recall assay, presenting altered baseline freezing profiles and a significantly reduced association index (AI) relative to controls (**Fig. 1g’’**). Collectively, these behavioral data establish a progressive timeline of cognitive dysfunction that manifests after 10 weeks but not after 5 weeks of the HF/HBP regimen.

### Early Systemic Vascular Stress Induces Region-Specific Myelin Thinning and Oligodendroglial Disruption

Myelination is essential for neural circuit plasticity and memory consolidation, with clinical evidence demonstrating white matter structural remodeling following complex task learning ^12–14^. Conversely, chronic systemic hypertension is a primary driver of clinically apparent white matter abnormalities^15^, where cognitive decline frequently correlates with profound demyelination^16,17^. To determine whether the HF/HBP model recapitulates these early structural alterations, we evaluated myelin architecture and oligodendroglial cell densities at the 5-week mark across two prominent white matter tracts: the corpus callosum and cerebellum.

Ultrastructural analysis via transmission electron microscopy (TEM) revealed that 5 weeks of the HF/HBP regimen significantly increased the g-ratio within the corpus callosum, indicating a reduction in myelin sheath thickness (**Fig. 2a,b**). To determine whether this thinning was restricted to specific axon calibers, we stratified the axonal population into four size-based quartiles (Q1 representing the smallest 25% of axons and Q4 representing the largest 25%). The g-ratio elevation remained consistent across all quartiles, demonstrating a uniform, size-independent thinning of the myelin sheath (**Fig. 2b’**).

**Figure 2.**
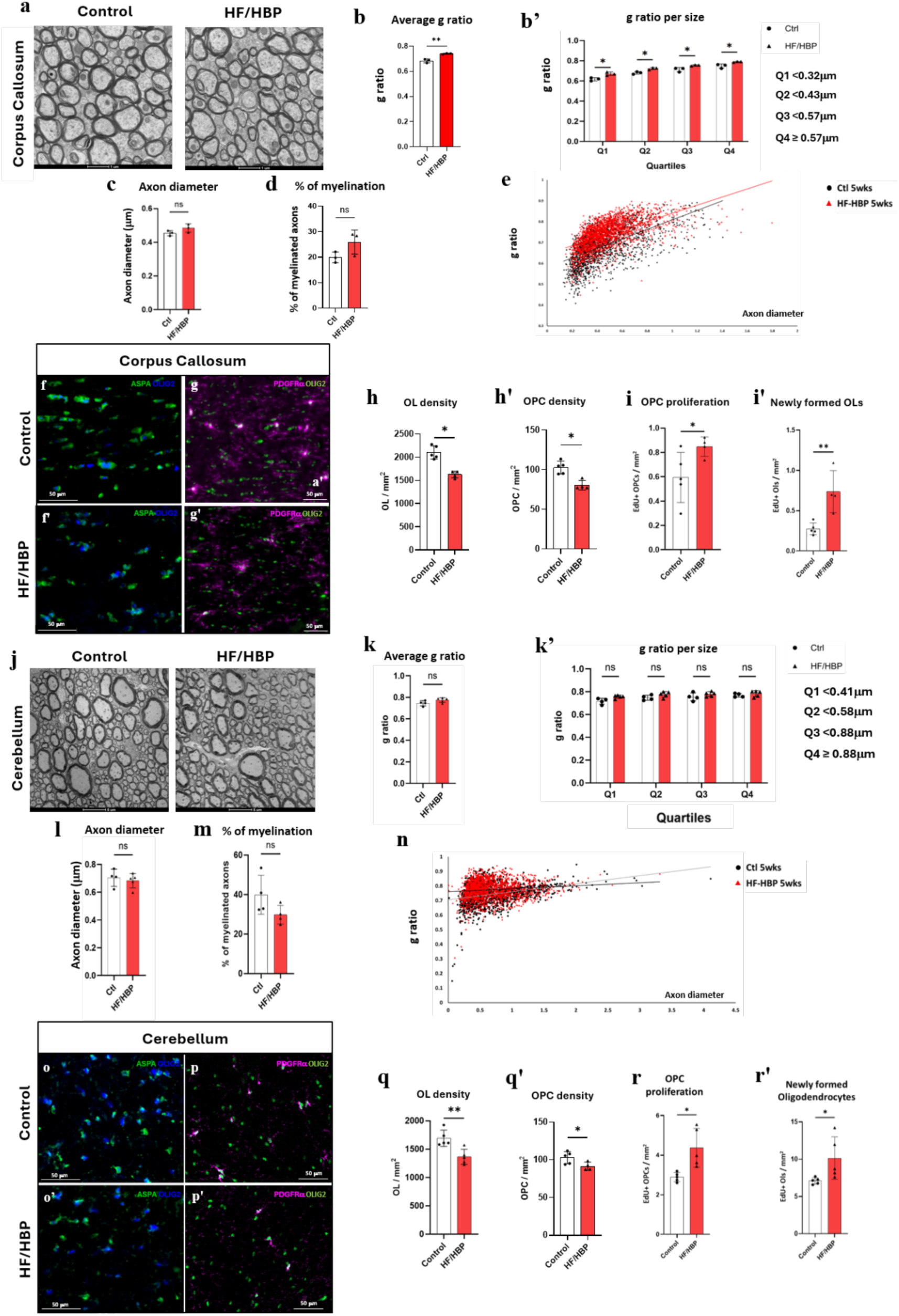
HF/HBP Diet Reduces Corpus Callosum Myelin Thickness and Oligodendroglial densities after 5 weeks of diet. (**a**) Representative corpus callosum electron microscopy pictures. (**b**) Corpus callosum’s axons g ratio (n=3 Ctl, 3 HF/HBP; Mann-Whitney, p=0.050). (**b’**) g ratio per size quartile (2way ANOVA, Diet effect p = 0.014; Multiple comparisons: Q1 p = 0.001, Q2 p = 0.009, Q3 p = 0.010 and Q4 p = 0.011). (**c**) Corpus Callosum’s axon diameter (Mann-Whitney: p = 0.200). (**d**) Corpus Callosum’s average percentage of myelinated axons (Mann-Whitney: p = 0.200). (**e**) Corpus callosum g ratio / axon diameter curves with linear regression of control and HF/HBP axons (ANCOVA: p < 0.001). (**f**) Representative pictures of corpus callosum oligodendrocytes (ASPA+/OLIG2+cells) at 5 weeks in Control or (**f’**) HF/HBP mice. (**g**) Representative pictures of corpus callosum OPCs (PDGFRα+/OLIG2+ cells) in 5 weeks Control or (**g’**) HF/HBP mice. (**h**) Corpus callosum oligodendrocyte (n=5 Ctl, 4 HF/HBP, Mann-Whitney: p = 0.016) and (**h’**) OPC density (Mann-Whitney: p = 0.016). (**i**) Corpus callosum proliferating OPCs (PDGFRα+/OLIG2+/EdU+) (Mann-Whitney: p = 0.008) and (**i’**) newly formed oligodendrocyte (PDGFRα-/OLIG2+/EdU+) density (Mann-Whitney: p = 0.016). (**j**) Representative cerebellum electron microscopy pictures. (**k**) Cerebellum average g ratio (n=4 Ctl, 4 HF/HBP, Mann-Whitney: P=0.413) and (**k’**) g ratio per axon diameter quartile (2way ANOVA: Diet p = 0.097; Multiple comparisons: Q1 p = 0.164, Q2 p = 0.363, Q3 p = 0.715 and Q4 p = 0.833). (**l**) Cerebellum’s axon diameter (Mann-Whitney: P = 0.591). (**m**) Cerebellum percentage of myelinated axons (Mann-Whitney: p = 0.114). (**n**) Cerebellum g ratio / axon diameter curves with linear regression (ANCOVA: p < 0.001). (**o**) Representative pictures of cerebellum oligodendrocytes in Control or (**o’**) HF/HBP male mice. (**p**) Representative pictures of cerebellum OPCs in Control or (**p’**) HF/HBP mice. (**q**) Oligodendrocytes (n=5 Control, 4 HF/HBP, Mann-Whitney: p = 0.008) and (**q’**) OPC density (Mann-Whitney: p = 0.032). (**r**) Cerebellum proliferating OPC (Mann-Whitney: P = 0.032) and (r’) newly formed oligodendrocyte (Mann-Whitney: p = 0.016) density during the 5^th^ week of diet

This myelin thinning occurred in the absence of gross structural loss, as the mean axonal diameter and overall percentage of myelinated fibers remained unaltered (**Fig.2c,d**). However, linear regression analysis of the g-ratio versus axon diameter slopes revealed a significant divergence between control and HF/HBP curves, confirming widespread, generalized myelin thinning (**Fig. 2e**).

To investigate the cellular dynamics underlying this myelin pathology, we used immunohistochemistry (IHC) to quantify mature oligodendrocytes and oligodendrocyte precursor cells (OPCs). We observed a 23% reduction in mature oligodendrocyte density and a 22% depletion of the OPC density within the corpus callosum of HF/HBP mice (**Fig. 2 f–h’**). Interestingly, this cellular loss triggered an endogenous regenerative response during the final week of the diet, characterized by a 51% increase in OPC proliferation and a 43% increase in the generation of newly differentiated oligodendrocytes (**Fig. 2i,i’**).

Intriguingly, the HF/HBP diet affected the cerebellum through a distinct pathological profile. Unlike the corpus callosum, the average cerebellar myelin g-ratio remained stable (**Fig. 2j–k’**), with no alterations detected in mean axon diameter or myelinated fiber percentages (**Fig. 2l, m**). Nevertheless, we observed a significant difference between the linear regression slopes of the control and HF/HBP cohorts, suggesting a subtle, subpopulation-specific effect on selected axonal diameters (**Fig. 2n**).

At the cellular level, the cerebellum closely mirrored the corpus callosum, displaying a 19% loss of mature oligodendrocyte density and an 11% reduction in OPC density (**Fig. 2o–q’**). This deficit similarly provoked homeostatic cell division and differentiation, yielding a 43% increase in OPC proliferation and a striking 168% spike in new oligodendrocyte generation (**Fig. 2r,r’**). In contrast to these cranial white matter tracts, the spinal cord exhibited complete protection, showing no alterations in either myelin architecture (**Fig. S1a–e**) or oligodendroglial populations (**Fig. S1f–h’**), underscoring a distinct spatial vulnerability to HF/HBP-induced demyelination.

We also investigated the extent to which sexual dimorphism affects the impact of the HF/HBP diet. While female HF/HBP mice exhibited significant weight gain relative to female controls, their overall metabolic trajectory was blunted compared to males (**Fig. S2a,b**), consistent with prior findings^18^. Cellular profiling of the female corpus callosum revealed an isolated reduction in OPC density, with preserved mature oligodendrocyte pools (**Fig. S2c–e**), whereas the cerebellum displayed the inverse phenotype: a reduction in mature oligodendrocyte density without depletion of the baseline OPC pool (**Fig. S2f–h**).

Furthermore, TEM analysis showed intact myelin sheaths within the female corpus callosum (**Fig. S2i–l**), although a significant shift in linear regression slopes suggested an initial, minor impact on a discrete subpopulation of axons (**Fig. S2m**). Because cerebellar and spinal cord myelin remained largely uninjured in males, these regions were not quantified in the female cohorts. In alignment with established literature demonstrating female mice resistance to early vascular and metabolic insults^19–21^ females presented with a substantially attenuated phenotype at 5 weeks. Consequently, only male mice were used for the remainder of the study.

### The HF/HBP Diet Selectively Activates the ISR across CNS Regions

The protein kinase R-like endoplasmic reticulum kinase (PERK) and downstream ISR pathway have been implicated in various, often contradictory, roles within the oligodendroglial lineage, depending on the pathological context^22–25^. To elucidate the molecular mechanisms driving the observed oligodendroglial defects in mice treated with the HF/HBP paradigm, we evaluated key proteomic indicators of ISR activation across distinct CNS regions.

In the corpus callosum of mice subjected to 5 weeks of the HF/HBP regimen, we detected a 30% increase in total PERK expression levels. In contrast, the protected HF/HBP cerebellum exhibited a decrease in PERK expression, while the spinal cord showed no baseline variation in total PERK levels (**Fig. 3a,b**).

**Figure 3.**
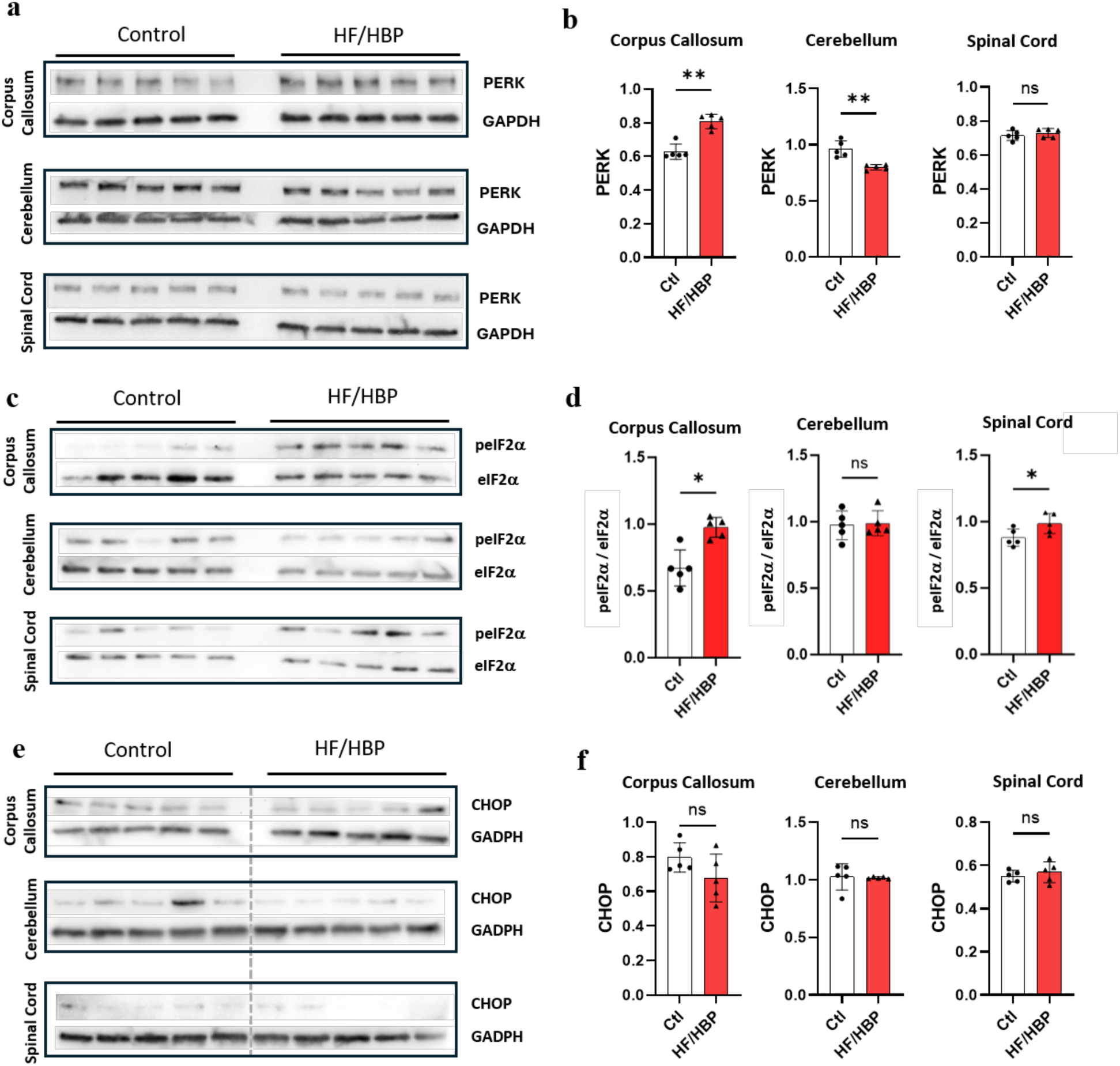
ISR Is Selectively Activated via PERK Upregulation in the Corpus Callosum but not in the Cerebellum after 5 weeks of HF/HBP Diet. (**a**) PERK western blots of 5 weeks Control and HF/HBP mice’s corpus callosum, cerebellum and spinal cord. GAPDH was used as a loading control. (**b**) PERK levels in 5 weeks Control and HF/HBP mice (n=5 Ctl and 5 HF/HBP, Mann-Whitney: corpus callosum p = 0.008; cerebellum p = 0.008; spinal cord p = 0.841). (**c**) p-eIF2α and total eIF2α western blots in 5 weeks Control and HF/HBP mice’s corpus callosum, cerebellum, and spinal cord. (**d**) peIF2α/eIF2α ratio in 5 weeks Control and HF/HBP mice (n=5 Ctl and 5 HF/HBP, Mann-Whitney test: corpus callosum p = 0.016; cerebellum p > 0.999; spinal cord p = 0.028). (**e**) CHOP western blots in 5 weeks Control and HF/HBP mice’s corpus callosum, cerebellum and spinal cord. (**f**) CHOP levels in 5 weeks Control and HF/HBP mice (n=5 Ctl and 5 HF/HBP, Mann-Whitney: corpus callosum p = 0.310, cerebellum p = 0.151, spinal cord p = 0.548)

Activated PERK kinase phosphorylates the translation initiation factor eIF2α^26^, resulting in reduced global protein synthesis^27,28^. Because the ratio of phosphorylated eIF2α to total eIF2α serves as a hallmark of functional ISR activation^29^, we quantified these isoforms using western blotting. Following 5 weeks of the dietary regimen, HF/HBP male mice exhibited a clear elevation in eIF2α phosphorylation within both the corpus callosum and spinal cord; however, no such increase was observed in the cerebellum (**Fig. 3c,d**).

Chronic or severe ISR activation is typically regulated by an endogenous negative feedback loop. When this homeostatic control fails, the downstream transcription factor C/EBP homologous protein (CHOP) accumulates in the nucleus, initiating an apoptotic cascade^30^. Intriguingly, no significant accumulation of CHOP was detected across any of the evaluated CNS tissues in HF/HBP mice (**Fig. 3e,f**).

### Pharmacological ISR Inhibition via 2BAct Rescues Myelin and Oligodendroglial Abnormalities

To examine how the observed ISR activation contributes to pathogenesis in the HF/HBP mouse model, we pharmacologically inhibited the pathway using 2BAct. This brain-penetrant, orally bioavailable small molecule activates eIF2B, thereby counteracting elevated levels of phosphorylated eIF2α. 2BAct has previously demonstrated therapeutic efficacy in normalizing chronic ISR activation^31,32^, preventing white matter lesion development, and preserving oligodendroglial integrity in genetic models of leukodystrophy^25^.

Mice were administered 2BAct in a control or high-fat diet and compared directly against untreated control or HF/HBP controls. To isolate the effects of metabolic variables, control cohorts were maintained on a glucose-matched control diet, with or without 2BAct supplementation, for 5 weeks (**Fig. 4a**). Pharmacological treatment did not alter gross metabolic trajectories; both control and HF/HBP mice receiving 2BAct gained equivalent weight compared to their respective untreated counterparts (**Fig. 4b**).

**Figure 4.**
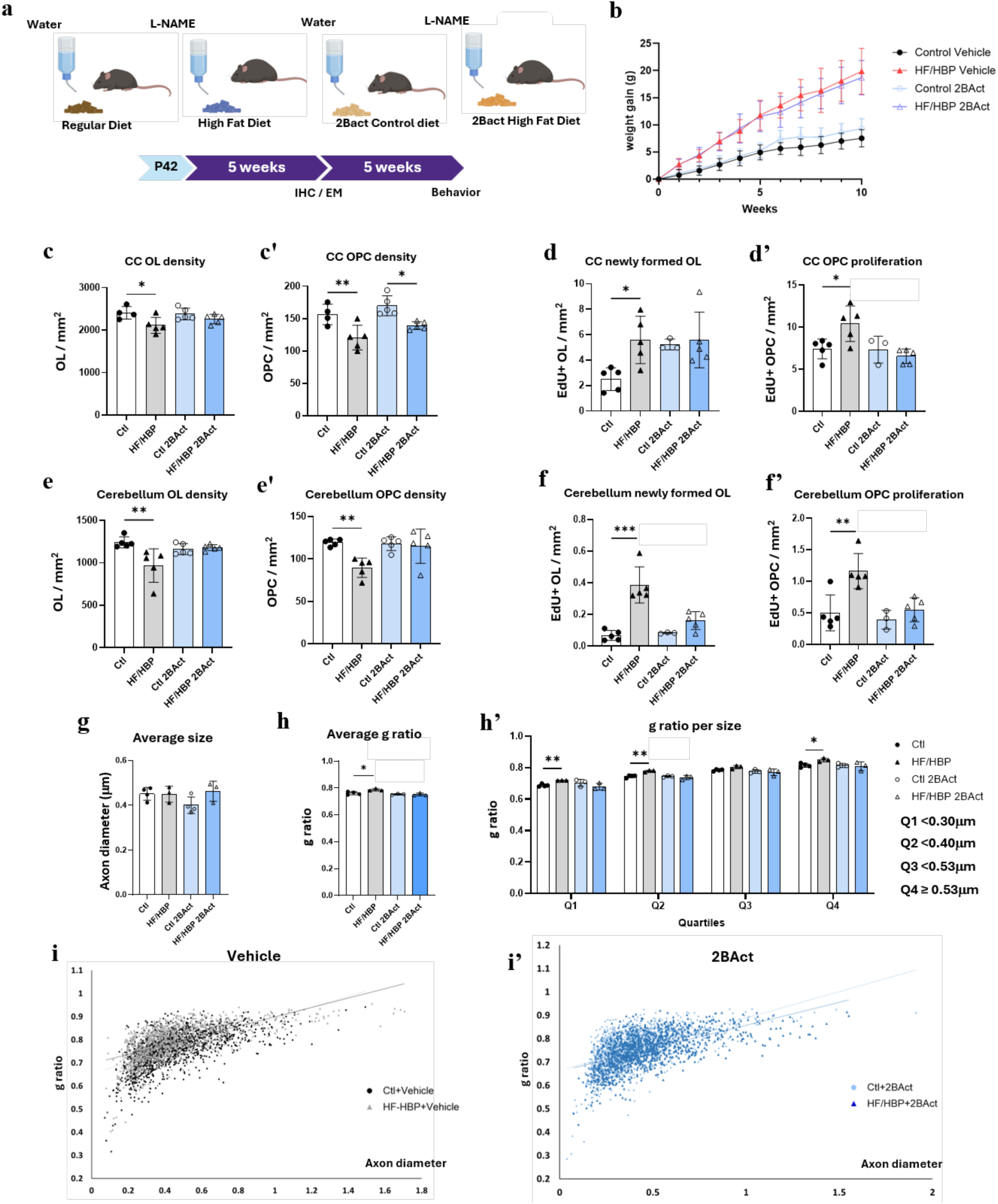
Drug Mediated ISR Inhibition with 2BAct Protects the Brain from HF/HBP Diet. (**a**) Experiment timeline. (**b**) Weight gain curves in mice treated with or without 2BAct (n=32 non treated (nt) Ctl; 33 nt HF/HBP; 32 2BAct Ctl; 33 2BAct HF/HBP, AUC analysis: nt-Ctl area p=42.34, Confidence Interval: 36.70-47.97; nt-HF/HBP area=108.1, Confidence Interval: 96.50-119.8; 2BAct-Ctl area=51.47, Confidence Interval: 45.07-57.86; 2BAct-HF/HBP area=103.8, Confidence Interval: 93.02-114.6). (**c**) Corpus callosum OLs (ASPA+/OLIG2+) (n=4 nt-Ctl; 5 nt-HF/HBP; 5 2BAct Ctl; 5 2BAct HF/HBP; 1way ANOVA p=0.032; nt Ctl vs. HF/HBP p = 0.020, 2BAct Ctl vs. HF/HBP p = 0.398) and (**c’**) OPCs density (PDGFRα+/OLIG2+) (1way ANOVA P < 0.001; nt Ctl vs. HF/HBP p = 0.005, 2BAct Ctl vs. HF/HBP p=0.010). (**d**) Corpus callosum PDGFRα-/OLIG2+/EdU+ newly formed OLs (n=5 nt Ctl; 5 nt HF/HBP; 3 2BAct Ctl; 5 2BAct HF/HBP; 1way ANOVA p=0.026; nt Ctl vs. HF/HBP p = 0.018, 2BAct Ctl vs. HF/HBP p = 0.943) and (**d’**) PDGFRα+/OLIG2+/EdU+ OPC density (1way ANOVA p = 0.006; nt Ctl vs. HF/HBP p = 0.014, 2BAct Ctl vs. HF/HBP p = 0.735). (**e**) Quantification of cerebellum ASPA+/OLIG2+ OL (n=5 nt Ctl; 5 nt HF/HBP; 3 2BAct Ctl; 5 2BAct HF/HBP; 1way ANOVA p = 0.008; nt Ctl vs. HF/HBP p = 0.003, 2BAct Ctl vs. HF/HBP p = 0.965) and (**e’**) PDGFRα+/OLIG2+ OPC density (1way ANOVA p = 0.005; nt Ctl vs. HF/HBP p = 0.003, 2BAct Ctl vs. HF/HBP p = 0.916). (**f**) Quantification of cerebellum PDGFRα-/OLIG2+/EdU+ OL (n=5 nt Ctl; 5 nt HF/HBP; 3 2BAct Ctl; 5 2BAct HF/HBP; 1way ANOVA; p < 0.001; nt Ctl vs. HF/HBP p < 0.001, 2BAct Ctl vs. HF/HBP p = 0.272) and (**f’**) PDGFRα+/OLIG2+/EdU+ OPC density (1way ANOVA p = 0.001; nt Ctl vs. HF/HBP p = 0.001, 2BAct Ctl vs. HF/HBP p = 0.630). (**g**) Average axon diameter in the corpus callosum (n=4 nt Ctl; 3 nt HF/HBP; 3 2BAct Ctl; 3 2BAct HF/HBP, 1way ANOVA p = 0.157; nt Ctl vs. HF/HBP p = 0.997, 2BAct Ctl vs. HF/HBP p = 0.097). (**h**) Average g ratio in the corpus callosum (n=4 nt Ctl; 3 nt HF/HBP; 3 2BAct Ctl; 3 2BAct HF/HBP; 1way ANOVA p = 0.004; nt Ctl vs. HF/HBP p = 0.012, 2BAct Ctl vs. HF/HBP p = 0.961). (**h’**) g ratio per axon diameter quartiles (2way ANOVA: Diet effect p = 0.007; Q1 nt Ctl vs HF/HBP p=0.027, Q1 2BAct Ctl vs HF/HBP p = 0.888, Q2 nt Ctl vs HF/HBP p = 0.010, Q2 2BAct Ctl vs HF/HBP p = 0.973, Q3 nt Ctl vs HF/HBP p = 0.342, Q3 2BAct Ctl vs HF/HBP P > 0.999, Q4 nt Ctl vs HF/HBP P = 0.088, Q4 2BAct Ctl vs HF/HBP p > 0.999). (**i**) Corpus callosum (n=4 nt Ctl; 3 nt HF/HBP; 3 2BAct Ctl; 3 2BAct HF/HBP) g ratio/axon diameter curves from nt (ANCOVA, p < 0.001) or (**i’**) 2BAct Control and HF/HBP mice (ANCOVA, p < 0.001)

At the cellular level, 2BAct treatment conferred robust protection within the white matter microenvironment. Following 5 weeks of the dietary regimen, 2BAct-treated HF/HBP mice were entirely protected against the decrease in oligodendrocyte density observed in untreated pathological conditions, displaying normal mature oligodendrocyte and OPC densities within the corpus callosum (**Fig. 4c,c’**). Intriguingly, both control and HF/HBP mice treated with 2BAct maintained elevated rates of mature oligodendrocyte generation, mirroring the endogenous regenerative response observed in untreated HF/HBP mice. However, 2BAct treatment completely suppressed the compensatory spike in OPC proliferation observed in untreated diseased animals (**Fig. 4d,d’**). Within the cerebellum, 2BAct similarly rescued oligodendroglial cell density (**Fig. 4e,e’**) and restored normal baseline rates of both oligodendrocyte differentiation and OPC proliferation (**Fig. 4f,f’**), validating its target engagement and survival benefits across distinct brain regions.

Ultrastructural analysis further confirmed that 2BAct treatment preserved structural myelin architecture. In the corpus callosum, small-molecule treatment successfully restored normal global g-ratios in HF/HBP mice without affecting mean axonal diameters (**Fig. 4g, h**). A quartile-based analysis demonstrated an absence of myelin thinning across all axonal calibers in the 2BAct-treated HF/HBP groups relative to the treated controls (**Fig. 4h’**). Despite this structural protection, linear regression analysis of g-ratio curves revealed that slopes from the 2BAct-treated control and HF/HBP cohorts remained significantly distinct, indicating that pharmacological provided only partial ultrastructural rescue (**Fig. 4i,i’**). Our data suggests that systemic pharmacological inhibition of the ISR via 2BAct helps oligodendrocytes to resist HF/HBP induced damages.

### Oligodendrocyte-Specific Perk Deletion Rescues HF/HBP-Induced Myelin Pathology and Cognitive Impairment

Given the beneficial effects of ISR inhibition on HF/HBP-induced pathogenesis, we investigated the impact of a dampened ISR response specifically in oligodendrocytes. To do this, we conditionally deleted the gene encoding the ISR-activating kinase PERK using *Plp1-CreER^T^; Perk^ff/ff^* mice. Following pre-diet administration of 4-hydroxytamoxifen to drive the *Perk^ff/ff^* mutation within mature oligodendrocytes, generally leading to an over 70% recombination^33–36^, mice were placed on either a control or HF/HBP diet for 5 or 10 weeks (**Fig. 5a**). Both mutant (Cre+) and control (Cre-) littermates displayed equivalent weight gain trajectories under their respective dietary conditions, confirming that the genetic deletion did not alter gross metabolic phenotypes (**Fig. 5b**).

**Figure 5.**
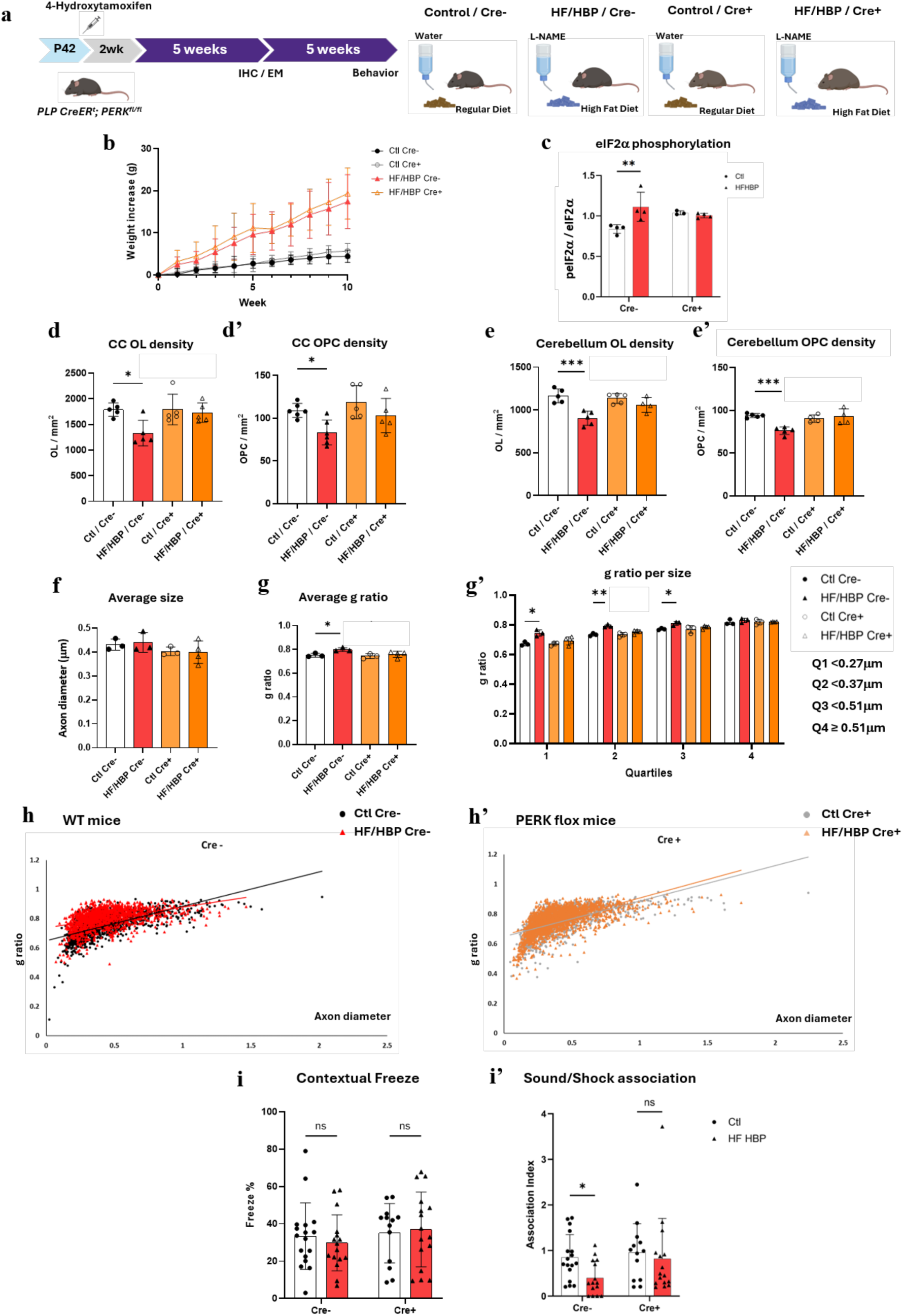
OL-Specific PERK Knockdown Fully Rescues HF/HBP-Induced Deficits. **(a)** Experiment timeline. **(b)** Weekly weight gain during 10 weeks of diet in mutant (Cre+) and non-mutant (Cre-) control or HF/HBP mice (n=21 Cre-Ctl; 21 Cre-HF/HBP; 15 Cre+ Ctl; 16 Cre+ HF/HBP, AUC: Cre-Control = 24.60, 95% Confidence Interval: 15.84-33.36; Cre-HF/HBP area = 87.50, 95% Confidence Interval: 68.51-106.5; Cre+ Control area = 27.72, 95% Confidence Interval: 22.77-32.67; Cre+ HF/HBP area = 90.50, 95% Confidence Interval: 74.60-106.4). **(c)** p-eIF2α/eIF2α ratio in Cre- or Cre+ 5 weeks Control or HF/HBP mice (n=4 Cre-Ctl; 4 Cre-HF/HBP; 3 Cre+ Ctl; 4 Cre+ HF/HBP, 2way ANOVA p = 0.042; Cre-Ctl vs HF/HBP p = 0.005; Cre+ Ctl vs HF/HBP p = 0.888). **(d)** Corpus callosum ASPA+/OLIG2+ OL (n=5 Cre-Ctl; 5 Cre-HF/HBP; 5 Cre+ Ctl; 5 Cre+ HF/HBP; 1way ANOVA p = 0.014; Cre-Ctl vs HF/HBP p = 0.011, Cre+ Ctl vs HF/HBP p = 0.896) and (**d’**) PDGFRα+/OLIG2+ OPC density (1way ANOVA p = 0.010; Cre-Ctl vs HF/HBP p = 0.023, Cre+ Ctl vs HF/HBP p = 0.250). **(e)** Cerebellum OL (1way ANOVA p < 0.001; Cre-Ctl vs HF/HBP p < 0.001, Cre+ Ctl vs HF/HBP p = 0.312) and **(e’)** OPC density (1way ANOVA p < 0.001; Cre-Ctl vs HF/HBP p < 0.001, Cre+ Ctl vs HF/HBP P = 0.811). **(f)** Corpus callosum’s axon diameter (n=3 Cre-Ctl; 3 Cre-HF/HBP; 3 Cre+ Ctl; 4 Cre+ HF/HBP; 1way ANOVA p = 0.431; Cre-Ctl vs HF/HBP p = 0.953, Cre+ Ctl vs HF/HBP p = 0.987). (**g**) Corpus Callosum’s average g ratio (1way ANOVA p = 0.034; Cre-Ctl vs. HF/HBP p = 0.036, Cre+ Ctl vs. HF/HBP p = 0.552) and (**g’**) g ratio per axon diameter quartile (2way ANOVA: diet p = 0.097; Q1 Cre-Ctl vs HF/HBP p = 0.038, Q1 Cre+ Ctl vs HF/HBP p = 0.588, Q2 Cre-Ctl vs HF/HBP p=0.006, Q2 Cre+ Ctl vs HF/HBP p = 0.391, Q3 Cre-Ctl vs HF/HBP p=0.056, Q3 Cre+ Ctl vs HF/HBP p = 0.780, Q4 Cre-Ctl vs HF/HBP p = 0.785, Q4 Cre+ Ctl vs HF/HBP p > 0.999). (**h**) Corpus callosum g ratio / axon diameter curves from Cre– (ANCOVA, p <0.001) or (**h’**) Cre+ Control and HF/HBP mice with linear regressions (ANCOVA: p = 0.2791). (**i**) 10 weeks Contextual Freeze (n=17 Cre-Ctl; 16 Cre-HF/HBP; 13 Cre+ Ctl; 16 Cre+ HF/HBP; Multiple Mann-Whitney: Cre-Ctl versus HF/HBP: p = 0.192; Cre+ Ctl versus HF/HBP: p = 0.619). (**i’**) 10 weeks’ AI (n=17 Cre-Ctl; 15 Cre-HF/HBP; 13 Cre+ Ctl; 16 Cre+ HF/HBP; Multiple Mann-Whitney: Cre-Ctl versus HF/HBP: p = 0.030; Cre+ Ctl versus HF/HBP: p = 0.268)

At the molecular level, western blot analysis confirmed the impact of our genetic targeting; unlike their Cre-counterparts, Cre+ mice completely lacked the characteristic elevation in eIF2α phosphorylation under HF/HBP conditions, establishing that oligodendrocyte-specific PERK deletion effectively prevents localized ISR activation (**Fig. 5c**). Cellular profiling via immunohistochemistry revealed that this molecular ablation translated to robust cytoprotection. While Cre-animals exhibited the expected drop in oligodendroglial numbers, Cre+ HF/HBP mice retained normal mature oligodendrocyte and OPC densities within both the corpus callosum and cerebellum (**Fig. 5d–e’**).

This localized cellular survival directly correlated with structural preservation of the myelin sheath. Ultrastructural examination of the corpus callosum in Cre+ HF/HBP mice revealed preserved myelin thickness, with normal global g-ratios and no alterations in baseline axon diameter or overall myelination percentages (**Fig. 5f,g**). Furthermore, a quartile-based caliber analysis confirmed that myelin sheaths were protected across all axon sizes (**Fig. 5g’**). Oligodendrocyte specific PERK KO fully protected myelin against the HF/HBP diet, as linear regression curves for Cre+ control and HF/HBP cohorts remained overlapping, contrasting with the divergent slopes seen in their Cre-littermates (**Fig. 5h,h’**).

Finally, we evaluated whether this protection of the white matter microenvironment at the cellular and structural levels would prevent late-stage cognitive decline. Indeed, ISR modulation has previously been shown to act as a molecular constraint on memory consolidation^37–40^. Following 10 weeks on the dietary regimen, while Cre-HF/HBP mice displayed significant learning impairments, Cre+ mutant mice were fully protected, showing no reduction in their AI during the auditory-cued recall assay (**Fig. 5i,i’**). Collectively, these findings demonstrate that cell-autonomous disruption of PERK signaling shields the oligodendroglial lineage from systemic vascular stress, preserving white matter architecture and preventing downstream cognitive deficits.

### ISR activation in human oligodendrocytes correlates with small vessel/cerebrovascular disease burden

To assess the relevance of these findings to human VaD, we examined the activation of the ISR in the white matter of the middle frontal gyrus from seven human brain autopsy donors spanning a range of small vessel/cerebrovascular disease burden with varying severities of VaD (ages 75-88 years, both sexes). Middle frontal cortex region sections stained with p-eIF2α immunohistochemistry (**Fig. 6a**). Hematoxylin and eosin (H&E) stained slides from middle frontal cortex were also examined by brightfield microscopy to identify white matter rarefaction (WMR), a common feature of VaD related to WMH^41^. The severity of small vessel disease (SVD) was evaluated using the SVD burden index previously described by Gonzalez et al.^42^, which harmonizes six National Alzheimer’s Coordinating Center (NACC) NP11 data from cerebrovascular domains — arteriolosclerosis, white matter rarefaction (WMR), cerebral amyloid angiopathy (CAA), microinfarcts, gross infarcts/lacunes, and microbleeds — onto 0-to-3 scales and sums them into a continuous SVD burden index ranging from 0 to 18. Briefly, and as previously described, for severity domains (arteriolosclerosis, WMR, CAA), National Alzheimer’s Coordinating Center (NACC) NP11 severity ratings were used directly, with ordinal descriptors (“none,” “mild,” “moderate,” “severe’) converted to numerical values 0–3. For count-based domains (microinfarcts, gross infarcts/lacunes, microbleeds), regional counts were summed across brain regions and binned as 0 = none, 1 = one lesion, 2 = two lesions, and 3 = three or more lesions. When at least five of the six domains were available, a scaled score was calculated as (sum of available domains / n present) × 6 to account for missing data. Domain scoring was performed independently of the immunofluorescence results. Tissue was also stained by immunofluorescence for the ISR marker p-eIF2α and the oligodendrocyte marker ASPA (**Fig. 6b**). Notably, we found a clear positive correlation between SVD burden and the percentage of p-eIF2α⁺/ASPA⁺ dual-positive cells in the human subcortical white matter, aligning with our findings in the murine HF/HBP model (**Fig. 6c**).

**Figure 6.**
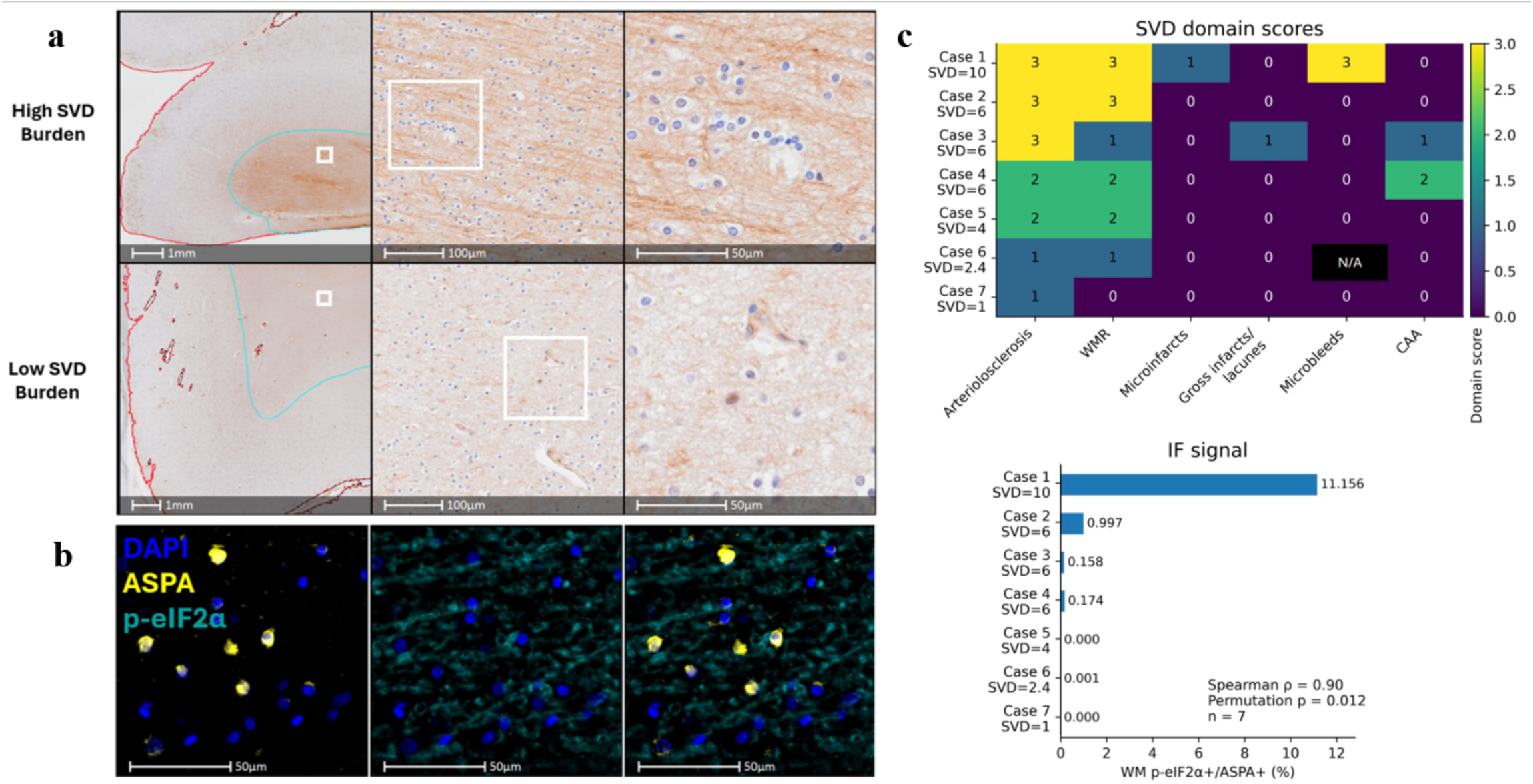
Human middle frontal gyrus white matter p-eIF2α signal increases with SVD burden. (**a**) Representative brightfield immunohistochemistry images show p-eIF2α staining in human middle frontal gyrus white matter from cases at high and low ends of the SVD burden range. (**b**) Representative multiplex immunofluorescence images of human middle frontal gyrus white matter showing p-eIF2α, aspartoacylase (ASPA), and DAPI, with merged signal shown where indicated. (**c**) Case-level heatmap of SVD domain scores and aligned WM p-eIF2α⁺/ASPA⁺ quantification in the human middle frontal gyrus cohort (n = 7). SVD domains were scored from 0 to 3 and included arteriolosclerosis, white matter rarefaction (WMR), microinfarcts, gross infarcts/lacunes, microbleeds, and cerebral amyloid angiopathy (CAA). Black/NA cell indicates missing domain-level data; the continuous SVD burden score was scaled to available domains, as described by Gonzalez et al^42^. The bar plot shows medium-threshold white matter p-eIF2α⁺/ASPA⁺ signal as a percentage of all nucleated cells in the region of interest. SVD burden association with p-eIF2α⁺/ASPA⁺ signal: Spearman rank correlation; ρ = 0.90; exact two-tailed permutation p = 0.012.

## Discussion

VaD is characterized by chronic memory impairment, executive dysfunction, and a correlation with WMH. Deciphering the molecular mechanisms that govern white matter vulnerability is a critical prerequisite for engineering targeted disease-modifying interventions. Despite standing as a leading global cause of cognitive decline^1^, the field has lacked non-surgical preclinical models that faithfully recapitulate both the cognitive decline and the underlying tissue damage seen in human VaD patients. In this study, we demonstrated that a non-surgical, dual-hit high-fat/high-blood-pressure (HF/HBP) mouse model effectively models these foundational clinical features.

In clinical settings, a definitive diagnosis of VaD requires evidence of multi-domain cognitive impairment, typically pairing memory deficits with decline in at least one other distinct cognitive parameter, such as agnosia or aphasia^3^. Here, we observed a progressive, multi-domain cognitive decline in the HF/HBP mice at the 10-week mark across three distinct behavioral paradigms. Spatial working memory was severely compromised in the novel object location task, while short-term recognition deficits were apparent in the novel object recognition test, where HF/HBP mice failed to distinguish object novelty. This inability to differentiate novel objects strongly mirrors the clinical presentation of visual agnosia observed in human VaD cohorts.

Furthermore, 10-week HF/HBP mice displayed marked neuropsychiatric-like alterations, manifesting as heightened thigmotaxis and diminished exploratory ambulation in the OF test, alongside exaggerated freezing behaviors during contextual fear conditioning. Interestingly, these behavioral phenotypes were absent at 5 weeks, despite being the earliest established time point for systemic cardiovascular dysfunction in this model^10^. This delayed manifestation aligns with the clinical progression of VaD, where systemic microvascular and metabolic insults accumulate subclinically long before transitioning into overt cognitive symptoms^43^.

Crucially, our data reveal that this late-stage behavioral decline is preceded by early, localized cellular and structural pathologies. At 5 weeks, well before the onset of behavioral deficits, HF/HBP mice exhibited a loss of oligodendroglial cells across the corpus callosum and cerebellum, alongside myelin thinning in the callosal tract. While magnetic resonance imaging remains the clinical gold standard for identifying WMH in a human pathological context, its spatial resolution is insufficient to detect early structural abnormalities. Using transmission electron microscopy, we captured these myelination defects at their onset.

Recent evidence indicates that subtle microenvironmental changes occur prior to visible WMH formation^44^. Because the HF/HBP model captures this pre-lesional window, it represents a powerful translational tool for evaluating early preventative and therapeutic interventions. Furthermore, the attenuated neuropathology observed in female mice at 5 weeks is consistent with reports demonstrating relative female resistance to early high-fat diet consumption^45^ and hypertensive stress^46^, driven in part by the vasoprotective effects of endogenous estrogens^19–21^.

The regional specificity of our model is further emphasized by the preservation of myelin architecture and oligodendroglial density within the spinal cord, indicating that the HF/HBP model selectively targets brain white matter. The spinal cord is spared in pure human VaD, further supporting the relevance of our HF/HBP model in mimicking this disorder.

The upstream molecular etiology of white matter defects in VaD remains elusive. Here, we provide evidence that ISR activation, which has been shown to play a critical role in the response to metabolic stress^47^, is a key factor in human VaD and in our murine model. The ISR pathway has been shown to exert divergent, context-dependent, effects within the oligodendroglial lineage. During neuroinflammatory insults, prolonged ISR activation in oligodendrocytes supports cell survival and promote remyelination^22,24^. Conversely, in models of intrinsic cell stress, such as Jimpy mutant mice, ISR inhibition dramatically enhances oligodendrocyte survival, myelination efficiency, and overall lifespan^25^. Chronic ISR activation in oligodendrocytes has been shown to result in myelin thinning^24^, similar to what is observed here in the HF/HBP model.

In the HF/HBP model, the expression levels of the ISR-triggering kinase PERK correlated with regional tissue vulnerability. Moreover, systemic pharmacological intervention via the eIF2B activator 2BAct provided protection to the mature oligodendrocyte pool, restored baseline oligodendroglial densities within the brain, and partially rescued callosal myelination. These findings align with a previous observation in a rat arterial occlusion model treated with the ISR inhibitor ISRIB^48^, which reversed neuropathological and behavioral aspects of the model.

Our genetic interventions utilizing *Plp1-CreER^T^; Perk^ff/ff^* mice provided mechanistic support for ISR involvement. First, genetic ablation of *Perk* in mature oligodendrocytes prevented the elevation of eIF2α phosphorylation under HF/HBP conditions. Because the brain expresses three other core eIF2α kinases capable of sensing distinct physiological stressors (HRI, GCN2, and PKR)^49^, this identifies PERK as the likely upstream kinase responsible for driving ISR activation during systemic metabolic and hypertensive stress. Second, the inactivation of *Perk* in mature oligodendrocytes provided protection to the myelinating cells and the myelin sheath, which correlated with the rescue of associative memory at 10 weeks. These findings underscore the pivotal role of oligodendrocytes in the initiation and progression of VaD.

Finally, we identified a direct association between SVD burden and ISR pathway activation in oligodendrocytes from human donor tissue. The favorable outcomes observed when modulating the ISR in our preclinical mouse HF/HBP model suggest that therapeutic strategies targeting this pathway could benefit patients with VaD. Although the ISR inhibitor 2BAct used in this study is not a clinical candidate due to off-target effects^32^, the CNS-penetrant ISR inhibitor Fosigotifator is currently in clinical trials for vanishing white matter disease (ClinicalTrials.gov Identifier: NCT05757141)^50^. This clinical precedent demonstrates the feasibility of translating a similar ISR-targeted approach to VaD.

## Limitations of the Study

We acknowledge several limitations within the present study. First, Schiattarella et al. previously noted a decrease in total running distance during exercise exhaustion assays in 5-week HF/HBP mice^10^. While this systematic physical vulnerability could introduce a confounding variable in behavioral tests that require ambulation, our internal velocity metrics demonstrated no significant differences in mean speeds across the OF, NOL and NOR tasks, suggesting that physical exhaustion did not bias our cognitive endpoints. Nevertheless, this factor could become confounding in analyses involving mice on a diet for extended durations.

Second, the long-term histopathological defects of female mice under chronic HF/HBP stress remain unknown. A comprehensive characterization of female neurovascular responses at late-stage time points is essential for future investigations. Finally, our analysis was restricted to three primary white matter structures (the corpus callosum, cerebellum, and spinal cord). A high-resolution, brain-wide mapping study correlating local ISR activation profiles with oligodendroglial damage would be highly valuable in determining whether other memory-critical networks exhibit similar vulnerabilities.

## Methods

### Animals

Mice were housed at the Center for Comparative Medicine of Northwestern University. We acquired postnatal day 35 (P35) wild-type C57BL/6J mice from Jackson Laboratory (strain #:000664) and allowed them to acclimatize to the facility for one week prior to initiating the dietary regimen. The *Plp1-CreER^T^; Perk^ff/ff^* strain was developed by crossing *B6.Cg-Tg(Plp1-cre/ERT)3Pop/J* strain we previously generated^33^ with *Eif2ak3^tm1.2Drc^/J* generously provided by Dr Doug Cavener^51^. *Plp1-CreER^T^; Perk^ff/ff^* mice were bred and maintained at the Northwestern University mouse facility. All animal studies adhered to the guidelines and were approved by the Institutional Animal Care and Use Committee of the Northwestern University. To minimize the number of mice used, behavioral cohorts were perfused for immunohistochemistry (IHC), electron microscopy (EM), or protein analysis 24 h after the final day of testing. For behavioral assays, mice were subjected to a basic human habituation protocol at the beginning of the diet: 1 minute of interaction with every mouse individually, then 5 minutes with the experimentator hand still in the cage. The individual interaction was reiterated every week during weight recording. Before testing, mice were allowed to acclimatize to the room for a minimum of 30 min at the beginning of every day. All mouse strains, equipment, and reagents are listed in the Key Resources Table.

### HF/HBP diet induction

At P42, mice were introduced to a new cage environment and assigned either a control diet (Teklad, 7912) or a high-fat diet (60 kcal% fat diet, Research Diet, D12492). Concurrently, they received either regular water or a 0.5 g/L solution of Nω-nitro-L-arginine methyl ester hydrochloride (L-NAME) dissolved in water, following the protocol outlined by Schiateralla et^10^. The diet and water were refreshed weekly, and the mice weights were recorded systematically. In the *Plp1-CreER^T^; Perk^ff/ff^* experimental group, mice were administered an intraperitoneal injection of 1 mg/day of 4-hydroxytamoxifen (4-OHT) at P42 for five consecutive days. A recovery period of 2 weeks was provided to mitigate stress before starting the dietary regimen at P63. Subsequently, the mice were housed in a new cage and sustained on the diet for 5 and 10 weeks prior to undergoing behavioral assessments or perfusion.

### 2BAct treatment

2BAct was incorporated into chow at a concentration of 300 mg/kg by Research Diet. To mitigate any potential effects arising from variations in sucrose levels between the control and experimental diets, a 10 kcal% fat diet was selected for the control mice, matching the sucrose content of the 60 kcal% fat diet provided to HF/HBP mice. Mice were introduced to the diet in a new cage at P42 and maintained on this diet for 5 and 10 weeks before behavioral testing and perfusion.

#### Open Field (OF)

The Open Field (OF) assay evaluates anxiety-like behaviors and potential changes in locomotion. Mice were assessed after 5 or 10 weeks on a specific diet, with different cohorts used at each time point. The mice were placed in a 54.6 cm × 54.6 cm OF arena located in a soundproof closet, where they were permitted to explore freely for 5 min. During the trial, the mice’s movements were tracked and recorded using Actimetrics LimeLight 5 software. After each trial, the OF boxes were cleaned with C-DOX and wiped with a 70% ethanol solution to eliminate feces, urine, and odors. This procedure was consistently applied to all subsequent trials involving OF boxes.

##### Metrics

We systematically monitored the central position of the mouse body and partitioned the OF into a 5x5 grid of 10.92 cm × 10.92 cm squares. We quantified the percentage of time the mice spent in the central square (“% of time in center’) and in any of the four corner squares (“% of time in corner’), as well as the average speed of the mouse. Mice inherently exhibit aversion to open spaces and typically remain close to the walls. Consequently, an increase in the time spent in the corners and a decrease in the time spent exploring the center are indicative of heightened anxiety. The average speed provides insights into mouse locomotion; however, a reduced speed may also correlate with anxiety.

#### New Object Location (NOL)

##### Training Phase

Two hours following the OF test, the mice were trained for the NOL task. The mice were reintroduced into the OF arena, where identical objects were positioned equidistantly from both mice and the arena walls. The trial commenced with the mice facing the wall opposite their starting point, allowing them to explore the arena freely for 10 min. This exploratory behavior was recorded and tracked using LimeLight 5 software, and the time spent investigating each object was calculated.

##### NOL Test Phase

Two hours after training, the mice were reintroduced to an identical setup. The object located on the right was moved to a new location. The mice were allowed to explore the area independently for 10 min before being returned to their original cages for the night.

##### Metrics

The time spent within a 9 cm radius circle centered on the object was defined as “interaction time.” The Recognition Index (RI) was calculated using the following formula: RI = %Time exploring an object / % time exploring both objects. The RI determined during the training phase (RIspont) indicates the spontaneous preference of a mouse for one of the objects, with an RI_spont_ of 0.5 indicating no preference and an RI > 0.5 indicating a preference for the object on the right side. The RI calculated during the NOL test phase (RI_shifted_) was normalized by RI_spont_ to derive the normalized RI (RI_norm_), thereby mitigating the bias induced by any inherent preference of the mouse for one of the objects. An RI_norm_ of 1 indicates no preference for either object, whereas an RI_norm_ > 1 suggests increased interaction time with the shifted object, which is expected in normal mice. A decrease in RI_norm_ indicates impaired spatial memory.

#### New Object Recognition (NOR)

The next day, the mice were reintroduced to the OF boxes, where they encountered the same object on their left and a novel object on their right side. This arrangement mirrored the configuration used during the NOL training phase. We recorded interactions with both objects for 10 min using Limelight 5.

##### Metrics

The Recognition Index was calculated during the NOR test and normalized by the spontaneous RI (RI_spont_) to adjust for any inherent bias toward a specific object. A RI_norm_ of 1 indicates no preference for either object, whereas an RI_norm_ greater than 1 suggests increased interaction time with the novel object, which is anticipated in normal mice. A reduced RI_norm_ in the NOR is indicative of memory deficits and an impaired ability to discriminate between shapes, similar to agnosia observed in humans.

#### Fear Conditioning (FC)

##### FC training phase

Two hours after the NOR test, the mice were placed in a shock chamber located within a soundproof closet. We recorded the percentage of freezing for 10 min using Actimetrics FreezeFrame 5 software. During the trial, a 75 dB tone was played for 30 s, followed by a 0.7 mA electric shock delivered through the metallic bars of the shock cage. This sequence was repeated three times consecutively before the mice were returned to their original cages overnight. Each video was manually validated, and any mouse that did not receive three consecutive shocks was excluded from the study. Prior to introducing the mice into the room, a control run was conducted by playing the tone in each soundproof closet, and a sound level meter (RadioShack) was used to confirm that no sound was detectable from outside the soundproof closets. It was also ensured that each mouse received an identical shock, as the intensity of the shock is known to influence the mice’s memorization ^52^.

##### FC contextual test

The following day, the mice were placed back into a cleaned shock chamber and recorded for 5 min without administering a shock or sound. The percentage of freezing during the entire trial was calculated using FreezeFrame 5 software.

##### FC cued test

Two hours later, the mice were placed in a new box without metallic bars in a soundproof closet for 390 s. During this period, the same 75 dB tone used in the FC training phase was played once for 60 s without administering any shocks to the mice. The percentage of freezing during the trial was recorded using FreezeFrame 5 software. Subsequently, the mice were euthanized or perfused for electron microscopy (EM) or immunohistochemical (IHC) analysis.

##### Metrics

The percentage of freezing observed during the contextual test reflects the mouse’s ability to recall the context in which it received the aversive stimuli (shocks) and is thus termed “Contextual Freezing.” An increase in Contextual Freezing is often correlated with anxiety or heightened pain sensitivity. The percentage of freezing observed while the sound was played during the cued test is referred to as “Cued Freezing.” Given our interest in the mice’s ability to associate sound with a shock, we calculated the AI by normalizing Cued Freezing with Contextual Freezing to mitigate bias induced by anxiety. An AI greater than 1 indicates that the mouse exhibits more freezing behavior upon hearing the sound than when placed in the context where it was shocked, thereby demonstrating a learned association between sound and shock. A reduced AI suggests a low association between sound and shock, indicating diminished learning and memory.

#### Immunostaining

Mice were administered an intraperitoneal injection of 20 mL per 10 g of body weight of a sterile 2.5% Avertin (2,2,2-Tribromoethanol diluted in 2-Methyl-2-butanol) solution for anesthesia. They were then transcardially perfused with 1X PBS at 4 °C for 5 min to clear blood from the brain, then with a 4% paraformaldehyde (PFA) solution at 4 °C. The dissected brains and whole spines were post-fixed in 4% PFA at 4 °C overnight. The spinal cords were dissected the following day, and all samples were incubated in a 30% sucrose solution at 4 °C overnight. After a minimum of 24 h, the brains and spinal cords were embedded in O.C.T. compound for coronal sectioning using dry ice. The cerebellum was dissected and embedded separately for sagittal sections.

A minimum of six non-consecutive 10 µm sections were sliced onto glass slides using an NX70 cryostat (Epredia) and stored at -80 °C. After thawing the slides at room temperature (RT) for 30 min, epitope retrieval was conducted in a Citrate Buffer Solution at 90 °C for 30 min. The slides were then allowed to cool at RT for 30 min before quenching the PFA with a 10 mM Glycine TBS-T solution for one hour. Epitopes were blocked with a blocking buffer consisting of 5% bovine serum albumin (BSA), 1% 100X Triton, and 1% donkey serum in 1X PBS at RT for one hour. Subsequently, the slides were incubated with primary antibodies diluted in blocking buffer overnight at 4 °C. The following day, the slides were incubated for 1 h at RT with secondary antibodies diluted in blocking buffer. Slides were mounted using ProLongTM Gold Antifade Reagent with DAPI. Spinal cord images were acquired using a SlideView VS200 slide scanner, and other tissues were imaged using a ZEISS LSM 900 Airyscan2 confocal microscope. Slides were washed three times with PBS for 5 min at RT between each step.

To trace the proliferation and generation of new cells, the mice were intraperitoneally injected with 50 μL per 10 g of a 1 µg/mL 5-Ethynyl-2’-deoxyuridine (EdU) solution 7 days prior to perfusion. Following secondary antibody incubation, EdU was detected using the Click-iT Plus EdU Cell Proliferation Kit protocol (Thermo Fisher Scientific).

### Ultrastructural Analysis using Electron Microscopy

Mice were anesthetized using Avertin solution and subsequently perfused with 1X PBS for 5 min until brain blood clearance was achieved. This was followed by perfusion with an EM buffer at 4°C and pH 7.3, consisting of 0.1M Sodium Cacodylate buffer, 2.5% glutaraldehyde, and 4% paraformaldehyde (PFA). Fixed specimens underwent graded dehydration through a sequential ethanol series (30%, 50%, 70%, 90%, 95%, and 100%), followed by transitional clearing in propylene oxide. Dehydrated tissues were embedded in an Epon-812 resin mixture consisting of 44.63% EMBed-812, 33.44% DDSA, 20.25% NMA, and 1.68% BDMA. Resin polymerization was carried out at 60°C for 48 h. Observations were conducted using an FEI Tecnai Spirit G2 transmission electron microscope (TEM). The g ratio, defined as the ratio of the inner diameter to the outer diameter of a myelinated axon, was calculated for each axon within a 518.3504 mm² section (1900X magnification, with a minimum of 300 fibers per corpus callosum and 150 per cerebellum and spinal cord). The percentage of myelinated fibers was quantified using MyelTracer software from an average of at least ten images at 6800X magnification ^53^.

### Western Blot

Mice were perfused with freshly prepared PBS at 4°C, and the spinal cord and brain were promptly extracted and snap-frozen in liquid nitrogen. The corpus callosum and cerebellum were manually dissected on ice and immediately returned to dry ice. Proteins were extracted using RIPA buffer supplemented with protease and phosphatase inhibitors. Protein concentration was determined using the Pierce BCA Protein Assay. Samples were denatured with 200 mM dithiothreitol (DTT) loading dye and heat. A total of 25 µg of protein was loaded onto a 4–20% Mini-PROTEAN TGX Precast Protein Gels and electrophoresed at 70 V for 15 min, followed by 120 V for 45 min. Dry transfer was performed on a PVDF membrane, previously activated with methanol, or a nitrocellulose membrane using a Trans-Blot Turbo Transfer System (Bio-Rad). Membranes were blocked for 1 h at room temperature with 5% dry milk diluted in TBS-T solution and then incubated with primary antibodies overnight at 4°C. Membranes were then washed and incubated with secondary HRP-conjugated antibodies for 1 h at room temperature. Signal detection was performed using a ChemiDoc Touch Imaging System. Antibodies were stripped from the membranes using a stripping buffer, followed by incubation in 5% dry milk diluted in TBS-T, and then re-incubated with another antibody solution. The intensity of each band was measured using Fiji, normalized to the intensity of the surrounding background in the same column, and subsequently to the intensity of the normalized GAPDH loading control intensity.

### Postmortem Human Brain Analysis

The human middle frontal gyrus samples from 7 brains donors (Biggs Brain Bank, UT Health San Antonio) were used to create formalin-fixed paraffin embedded slides. All cases were scored using the Gonzalez et al. framework^42^, which harmonizes six NP11 cerebrovascular domains onto 0–3 scales: arteriolosclerosis, white matter rarefaction, cerebral amyloid angiopathy, microinfarcts, gross infarcts/lacunes, and microbleeds. The score is intended to capture cumulative vascular pathology across multiple lesion types and was shown to be associated with global cognition and odds of dementia. Immunohistochemistry & multiplexed immunofluorescence staining was performed on all cases using conventional methods, see Key Reagent table for antibodies and other reagents used in this study.

Whole slide images (WSIs) were acquired using an Olympus VS200 Virtual Slide System at 40X magnification with an Olympus XAPO objective. For the multiplexed IF stained sections, fluorophores were excited with a X-Cite NOVEM-IR (XT940) 9-channel LED fluorescence illuminator (Excelitas Technologies, Pittsburgh, Pennsylvania) and SpectraSplit® 7 (Kromnigon, Mölndal, Sweden) and seven channel filter set with the following exposure configuration: S-Split Blue (7.8 ms; DAPI), S-Split Red (31ms; Opal 620), S-Split Infra-red (1.49 s, Alexa 750). White matter regions of interest were segmented in HALO (v4.2.6399, Indica Labs), and multiplex IF cellular phenotypes were generated using the HALO HighPlex FL module. WM segmentations were manually QC’d and edited where needed, especially in high CeVD with tissue loss. Phenotypes were assigned to DAPI-positive, nuclear-expanded cell objects based on met/unmet fluorescence intensity thresholds for each channel. For the p-eIF2α channel, the medium-intensity threshold was used to identify p-eIF2α⁺/ASPA⁺ dual-positive cell objects, and the percentage was calculated using all nucleated cell objects in the white matter ROI as the denominator.

### Statistics

Statistical analyses and histograms were conducted using Prism-GraphPad software. A 95% confidence interval was employed for the area under the curve to assess differences in weight intake. The non-parametric unpaired Mann-Whitney test was used to compare the two conditions. For analyses involving more than two parameters, such as quartile and diet in the EM analysis across multiple groups, a two-way ANOVA with Šidák’s multiple comparisons was applied. When examining the effect of a single parameter, such as genotype or drug treatment, on multiple groups, a one-way ANOVA with Šidák correction was used to compare Control and HF/HBP diet mice within the same condition (Cre-/Cre+ or non-treated/2BAct). ANCOVA was used to compare the slopes of linear regressions between the two conditions. Given the distinct differences between Cre- and Cre+ *Plp1-CreER^T^; Perk^ff/ff^* and non-treated and 2BAct-treated mice, comparisons were restricted to the Control and HF/HBP within each condition using multiple unpaired Mann-Whitney tests. For human brain postmortem analysis, non-parametric Spearman rank correlation was used. Significance was assessed by exact two-tailed permutation across all 5,040 rearrangements of the IF values. Leave-one-out sensitivity analysis was performed by removing the highest-burden case and re-running the test.

## Acknowledgements

Behavioral assays were performed at the Northwestern University Behavioral Phenotyping Core Facility. Electron microscopy imaging was performed at the Northwestern University Center for Advanced Microscopy (RRID: SCR_020996), generously supported by NCI CCSG P30 CA060553 awarded to the Robert H Lurie Comprehensive Cancer Center using their FEI Spirit G2 TEM. We would like to thank Erdong Liu (Northwestern University), Gloria Wright (Northwestern University), Li Kai (Northwestern University), and Kat Gemperli (Northwestern University) for their critical assistance.

## Funding

This research was supported by NIH grants NIH/NINDS 5R01AG072080 (BP and MEF), R35NS137478 (BP), P30AG066546 (PI Seshadri; Neuropathology Core Leader: MEF), and the Dr. Miriam and Sheldon G. Adelson Medical Research Foundation (BP).

## Author contribution

EM, TG, WCC, MD, RK, IS, and MH performed the experiments. EM, TG, WCC, and MEF analyzed the data. EM, TG, FAS, EBT, MEF, and B.P. planned experiments. EM and B.P. wrote the manuscript with substantial contributions from all authors

## Competing interests

Authors declare that they have no competing interests.

## Materials & Correspondence

Any mice line or information required to reanalyze the data reported in this paper is available from Brian Popko upon request.

---

**Supplementary Figure 1.**
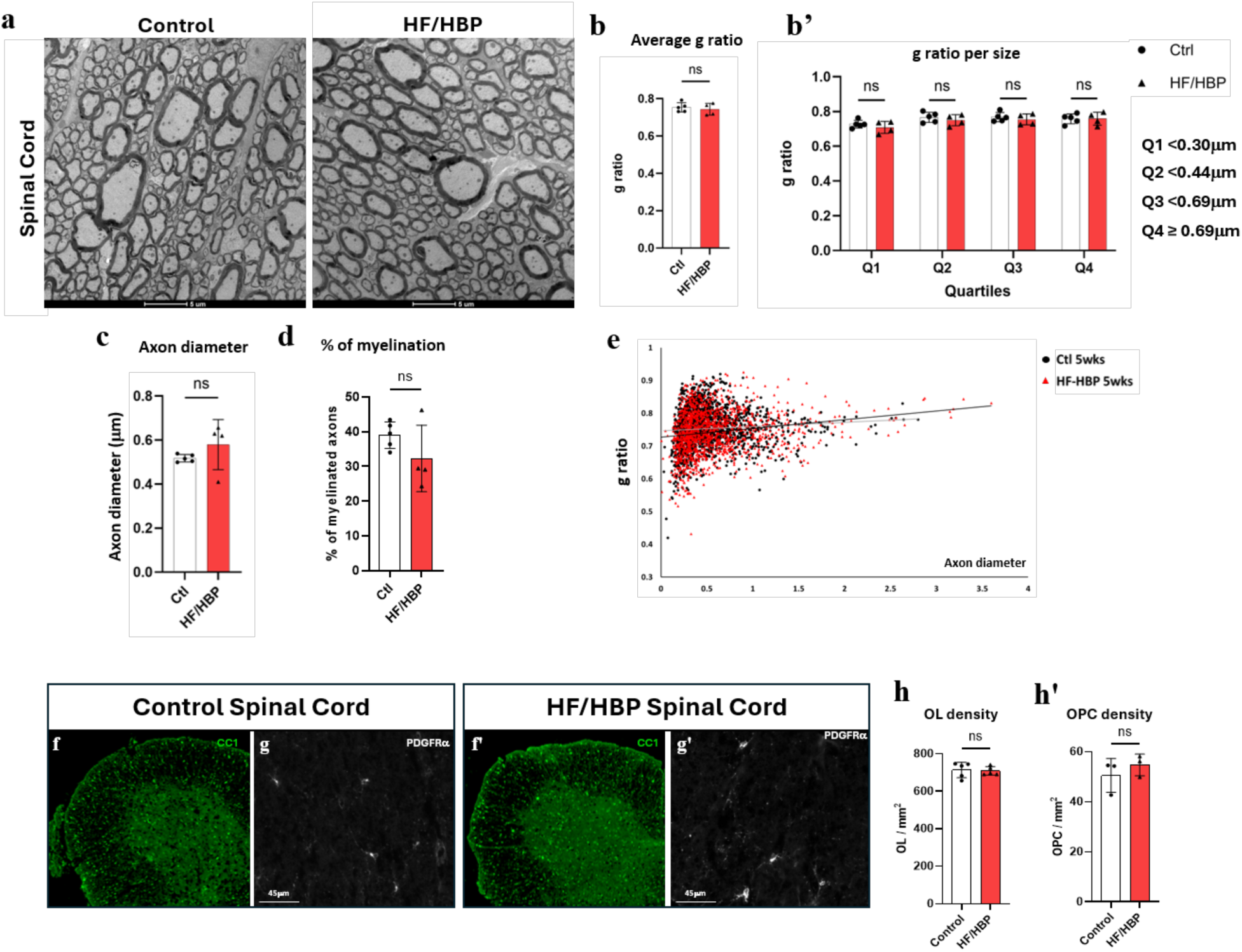
HF/HBP diet does not impact Spinal Cord White Matter. (**a**) Representative electron microscopic pictures of 5 weeks male spinal cords. (**b**) Spinal cord’s average g ratio (n=5 Ctl, 4 HF/HBP, Mann-Whitney, p = 0.730). (b’) Spinal cord’s g ratio per axon diameter quartile (2way ANOVA, diet p = 0.587; multiple comparisons: Q1 p = 0.873, Q2 p = 0.912, Q3 p = 0.951, and Q4 p > 0.999). (**c**) Spinal Cord average axon diameter (Mann-Whitney, p = 0.286). (**d**) Spinal’s cord average percentage of myelinated axons (Mann-Whitney, p = 0.286). (**e**) Spinal cord’s g ratio per axon diameter curves with linear regression (ANCOVA, p = 0.344). (**f-g**) Representative images of (**f**) spinal cord oligodendrocyte (CC1+) and (**g**) OPC (PDGFRα+) in 5weeks male mice. (**h**) Spinal cord’s oligodendrocyte (Mann-Whitney, p = 0.421) and (**h’**) OPC density (Mann-Whitney, p=0.200)

**Supplementary Figure 2.**
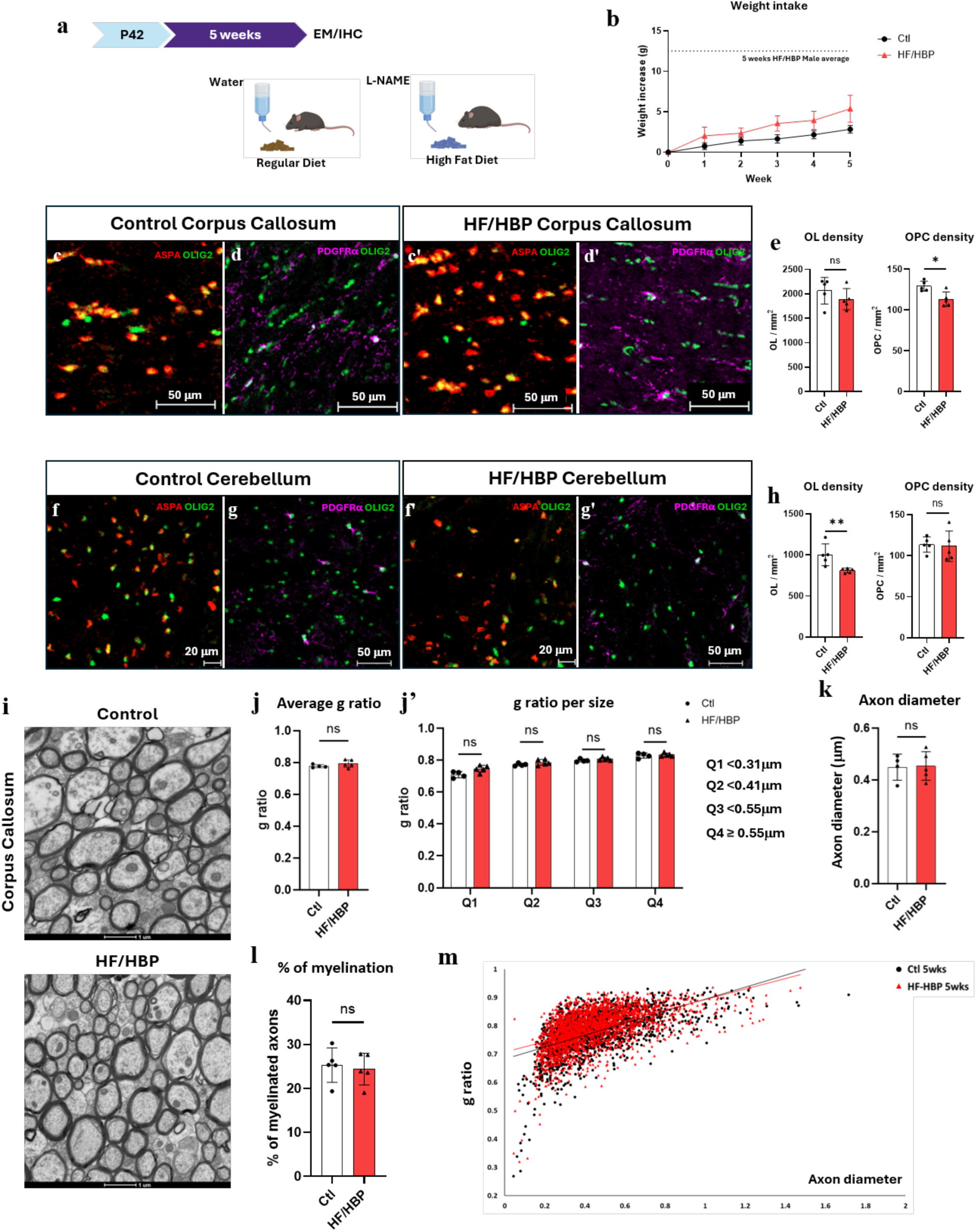
Female HF/HBP Mice exhibit a Weaker Phenotype. (**a**) Experiment’s timeline. (**b**) Weekly weight-gain curves of female mice (n=30 Ctl, 30 HF/HBP, AUC: Control area = 7.382, 95% Confidence Interval: 5.969-8.795; HF/HBP area = 14.58, 95% Confidence Interval:11.44-17.72). The dotted line represents the average 5 weeks weight intake in male HF/HBP diet mice (**c**) Representative pictures of corpus callosum oligodendrocytes (ASPA+/OLIG2+) in Control or (**c’**) HF/HBP mice. (**d**) Representative pictures of corpus callosum OPCs (PDGFRα+/OLIG2+) in Control or (**d’**) HF/HBP mice. (**e**) Oligodendrocyte and OPC density in 5weeks female mice corpus callosum (n= 5 Ctl and 5 HF/HBP; Mann-Whitney: oligodendrocyte p = 0.155; OPC p = 0.028). (**f**) Representative pictures of cerebellum oligodendrocytes in Control or (**f’**) HF/HBP mice. (**g**) Representative pictures of cerebellum OPCs in Control or (**g’**) HF/HBP mice. (**h**) Oligodendrocyte and OPC density in 5weeks female mice cerebellum (n=5 Ctl and 5 HF/HBP; Mann-Whitney: oligodendrocyte p = 0.004; OPC p = 0.345). (**i**) Representative electron microscopy pictures female mice’s corpus callosum. (**j**) Corpus Callosum’s axons average g ratio (n=4 Ctl and 5 HF/HBP; Mann-Whitney: p = 0.251). (**j’**) Corpus callosum’s axons g ratio per axon diameter quartile (2way ANOVA: diet p = 0.099; Multiple comparison: Q1 p = 0.108, Q2 p = 0.528, Q3 p = 0.528 and Q4 p = 0.982). (**k**) Corpus Callosum’s axon average axon diameter (Mann-Whitney: p = 0.900). (**l**) Corpus Callosum average percentage of myelinated axons (n=5 Ctl and 5 HF/HBP; Mann-Whitney: p = 0.709). (**m**) Female’s corpus Callosum’s g ratio / axon diameter curves with linear regression (n=4 Control and 5 HF/HBP; ANCOVA: p = 0.007)

## Key resources table

Reagents, resources, hardware and software used to conduct this study

| REAGENT OR RESSOURCE | SOURCE | IDENTIFIER |
| --- | --- | --- |
| **Chemicals and Ressources** |  |  |
| NFDM (Non-Fat Dry Milk Powder) / Skim Milk Powder | Bio World | 30620074 |
| AR2 Buffer | Biocare Medical | ORI6005T60 |
| 10x Tris/Glycine/SDS | BioRad | #1610732 |
| 2x Laemmli Sample Buffer | BioRad | #1610737 |
| 4–20% Mini-PROTEAN® TGX™ Precast Protein Gels | BioRad | #4561095 |
| Immun-Blot PVDF Membrane | BioRad | Cat #1620177 |
| Trans-Blot Turbo RTA Mini 0.2 µm Nitrocellulose Transfer Kit | BioRad | #1704270 |
| Precision Plus Protein™ Kaleidoscope™ Prestained Protein Standards | BioRad | #1610375 |
| Trans-Blot Turbo Mini Filter Paper | BioRad | #12023835 |
| 10X Phosphate-Buffered Saline, RNase-/DNase- and protease-free | Corning® | 46-013-CM |
| Decon's Pure Ethanol 200 Proof (24 x 1PINT) | Decon Labs | #2716 |
| Aqueous Glutaraldehyde EM Grade 25% | Electron Microscopy | #16210 |
| Benzyldimethylamine, (BDMA) | Electron Microscopy | #11400-25 |
| DDSA (Dodecyl Succinic Anhydride Specially Distilled) | Electron Microscopy | #13710 |
| Embed 812 Resin | Electron Microscopy | #14900 |
| NMA (Methyl-5-Norbomene-2, 3-Dicarboxylic Anhydride) | Electron Microscopy | #19000 |
| Osmium Tetroxide 2% Aqueous Solution | Electron Microscopy | #19152 |
| Paraformaldehyde, EM Grade, Purified | Electron Microscopy | #19200 |
| Propylen Oxide | Electron Microscopy | #20401 |
| Sodium Cacodylate Buffer, 0.2M, pH 7.4 | Electron Microscopy | #11652 |
| Sodium Hydroxide (Pellets/Certified ACS) | Fisher Chemical™ | S318-100 |
| 4-Hydroxytamoxifen ≥70% Z isomer (remainder primarily E-isomer) | Hello Bio | HB6040 |
| Click-iT™ Plus EdU Cell Proliferation Kit for Imaging, Alexa Fluor™ 594 dye | Invitrogen™ | C10639 |
| ProLong™ Gold Antifade Mountant with DNA Stain DAPI | Invitrogen™ | P36931 |
| Normal donkey serum | Jackson ImmunoResearch | Cat 017-000-121 |
| 2BAct | MedChemExpress | HY-125021 |
| DL-Dithiothreitol (Dtt) | Millipore Sigma | 43815 |
| Hydrochloric acid | Millipore Sigma | H9892 |
| Sucrose OmniPur® | Millipore Sigma | 8510-5KG |
| Rodent Diet With 10 kcal% Fat (Matching Sucrose to D12492) | Research Diets | D12450J |
| Rodent Diet With 10 kcal% Fat and 300mg/kg 2BAct | Research Diets | D24061001 |
| Rodent Diet With 60 kcal% Fat | Research Diets | D12492 |
| Rodent Diet With 60 kcal% Fat and 300mg/kg 2BAct | Research Diets | D24061002 |
| Glycine, Free Base | Research Products International | G36050-1000.0 |
| Sodium Phosphate Monobasic Monohydrate [Sodium dihydrogen phosphate monohydrate] | Research Products International | S23120-1000.0 |
| Tris Base Ultra Pure [Tris (Hydroxymethyl) Aminomethane] | Research Products International | T60040 |
| Tween 20 [Polyoxyethylenesorbitan Monolaurate] | Research Products International | P20370-0.5 |
| 2,2,2-Tribromoethanol | Sigma-Aldrich | T48402 |
| Bovine Serum Albumin | Sigma-Aldrich | A7906 |
| Citrate Buffer | Sigma-Aldrich | NC2081580 |
| Dimethyl sulfoxide | Sigma-Aldrich | D2650-100ML |
| Nω-Nitro-L-arginine methyl ester hydrochloride (L-NAME) | Sigma-Aldrich | N5751 |
| RIPA Buffer | Sigma-Aldrich | R0278 |
| Sunflower seed oil | Sigma-Aldrich | S5007 |
| 2-Methyl-2-butanol | Sigma-Aldrich | 152463 |
| Triton™ X-100 | Sigma-Aldrich | T8787 |
| EdU (5-ethynyl-2'-deoxyuridine) | Thermo Fisher Scientific | A10044 |
| DAB (3,3'-Diaminobenzidine) | Thermo Fisher Scientific | BDB550880 |
| Halt™ Protease and Phosphatase Inhibitor Cocktail | Thermo Fisher Scientific | 78442 |
| Methanol | Thermo Fisher Scientific | A412-4 |
| Tissue-Plus™ O.C.T. Compound | Thermo Fisher Scientific | 23-730-571 |
| ProLong™ Diamond Antifade Mountant | Thermo Fisher Scientific | P36970 |
| Pierce™ BCA Protein Assay Kits | Thermo Fisher Scientific | 23225 |
| Pierce™ Bovine Serum Albumin Standard Ampules, 2 mg/mL | Thermo Fisher Scientific | 23209 |
| Restore™ Western Blot Stripping Buffer | Thermo Fisher Scientific | 21063 |
| Sodium Hydroxide, 12.0 N, Ricca Chemical | Thermo Fisher Scientific | R74701201A |
| SuperSignal™ West Dura Extended Duration Substrate | Thermo Fisher Scientific | 34076 |
| **Antibodies** |  |  |
| Anti-mouse EIF2S1 (phospho S51) antibody [E90] (Rabbit) | Abcam | ab32157 |
| Antibody Diluent / Block 1X | Akoya Biosciences | ARD1001EA |
| AR6 Buffer | Akoya Biosciences | AR600250ML |
| Opal Anti-Ms + Rb HRP Kit | Akoya Biosciences | ARH3001KT |
| Opal 620 | Akoya Biosciences | FP1495001KT |
| Spectral DAPI | Akoya Biosciences | FP1490 |
| Mouse eIF2α Antibody (Rabbit) | Cell Signaling | Cat #9722 |
| Mouse GAPDH (14C10) Rabbit mAb #2118 | Cell Signaling | Cat #2118 |
| Mouse PERK (C33E10) Rabbit mAb | Cell Signaling | Cat #3192 |
| Rabbit Aspartoacylase antibody (ASPA) [N1C3-2] | GeneTex | GTX113389 |
| Donkey anti-Goat IgG (H+L) Cross-Adsorbed Secondary Antibody, Alexa Fluor™ 647 | Invitrogen | A-21447 |
| Donkey anti-Mouse IgG (H+L) Highly Cross-Adsorbed Secondary Antibody, Alexa Fluor™ 488 | Invitrogen | A-21202 |
| Donkey anti-Mouse IgG (H+L) Highly Cross-Adsorbed Secondary Antibody, Alexa Fluor™ 647 | Invitrogen | A-31571 |
| Donkey anti-Rabbit IgG (H+L) Highly Cross-Adsorbed Secondary Antibody, Alexa Fluor™ 488 | Invitrogen | A-21206 |
| Donkey anti-Rabbit IgG (H+L) Highly Cross-Adsorbed Secondary Antibody, Alexa Fluor™ 594 | Invitrogen | A-21207 |
| Goat anti-Rabbit IgG (H+L) Secondary Antibody, HRP | Invitrogen | Cat #31460 |
| Human p-eIF2α (pSer52) Rabbit Monoclonal Antibody | Invitrogen | 44-728G |
| Anti-mouse OLIG2 Antibody mouse monoclonal, 211F1.1 | Milipore Sigma | MABN50 |
| Mouse CHOP Polyclonal Antibody | Proteintech | 15204-1-AP |
| Mouse PDGF R alpha Antibody | R & D Systems | AF1062 |
| Anti-APC (Ab-7) Mouse mAb (CC-1) | Sigma-Aldrich | OP80 |
| Alexa Fluor™ 750 Goat Anti-Rabbit IgG H&L | Thermo Fisher Scientific | A-21039 |
| ImmPRESS® HRP Goat Anti-Rabbit IgG Polymer Detection Kit | Vector Laboratories | MP-7451-50 |
| **Experimental Models: Mice** |  |  |
| *C57BL/6J* | Jackson Laboratory | #000664 |
| *B6.Cg-Tg(Plp1-cre/ERT)3Pop/J* | Jackson Laboratory | DOI: 10.1002/gene.10154 |
| *Eif2ak3<sup>tm1.2Drc</sup>/J* | Provided by Dr D.Cavener | DOI: 10.1128/MCB.22.11.3864-3874.2002 |
| *PLP/CreER<sup>t</sup>; PERK<sup>fl/fl</sup>* |  |  |
| **Hardware** |  |  |
| FreezeFrame 5 | Actimetrics |  |
| Limelight 5 | Actimetrics |  |
| WaterMaze 6 | Actimetrics |  |
| PaperPal | CACTUS |  |
| PRISM-Graphpad | Dotmatics |  |
| HALO | Indica Labs |  |
| VS200 ASW | Olympus |  |
| FIJI | Open source |  |
| MyelTracer | Open source |  |
| BioRender | Science Suite Inc |  |
| Zen | Zeiss |  |
| **Hardware** |  |  |
| ChemiDoc™ Touch Imaging System | BioRad |  |
| Trans-Blot Turbo Transfer System | BioRad |  |
| Cryostar NX70 | Epredia |  |
| XCite-Xylis | Excelitas Technologies |  |
| XCite-NOVEM-IR | Excelitas Technologies | XT940 |
| Digital Camera C13440 | Hamatsu |  |
| Slideview VS200 (Slide Scanner) | Olympus |  |
| ZEISS LSM 900 Airyscan2 | Zeiss |  |
| Graphical tablet | HUION |  |
| Digital Sound Level meter | RadioShack | 33-2055 |
| FEI Tecnai G2 Spirit Twin TEM | Thermo Fisher Scientific |  |

## Notes

### Competing Interest Statement

The authors have declared no competing interest.

